# *In vivo* Evaluation and *in silico* Prediction of the Toxicity of Drepanoalpha^®^ hard capsules

**DOI:** 10.1101/2020.12.03.411124

**Authors:** Benjamin Z. Gbolo, K. N. Ngbolua, Damien S. T. Tshibangu, Patrick B. Memvanga, Dorothée D. Tshilanda, Aristote Matondo, Jason T. Kilembe, Bienvenu M. Lebwaze, Amandine Nachtergael, Pius T. Mpiana, Pierre Duez

**Affiliations:** Department of Biology, Faculty of Sciences, University of Kinshasa, P.O. Box 190, Kinshasa XI, Democratic Republic of the Congo.; Department of Chemistry, Faculty of Sciences, University of Kinshasa, P.O. Box 190 Kinshasa XI, Democratic Republic of the Congo.; Faculty of Pharmaceutical Sciences, University of Kinshasa, P.O.Box 212, Democratic Republic of the Congo; Department of Anatomopathology, Faculty of Medicine, University of Kinshasa, P.O.Box 212, Democratic Republic of the Congo; Unit of Therapeutic Chemistry and Pharmacognosy, Faculty of Medicine and Pharmacy, University of Mons (UMONS), 7000 Mons, Belgium

**Keywords:** Acute toxicity, Sub-acute toxicity, Drepanoalpha^®^ hard capsules

## Abstract

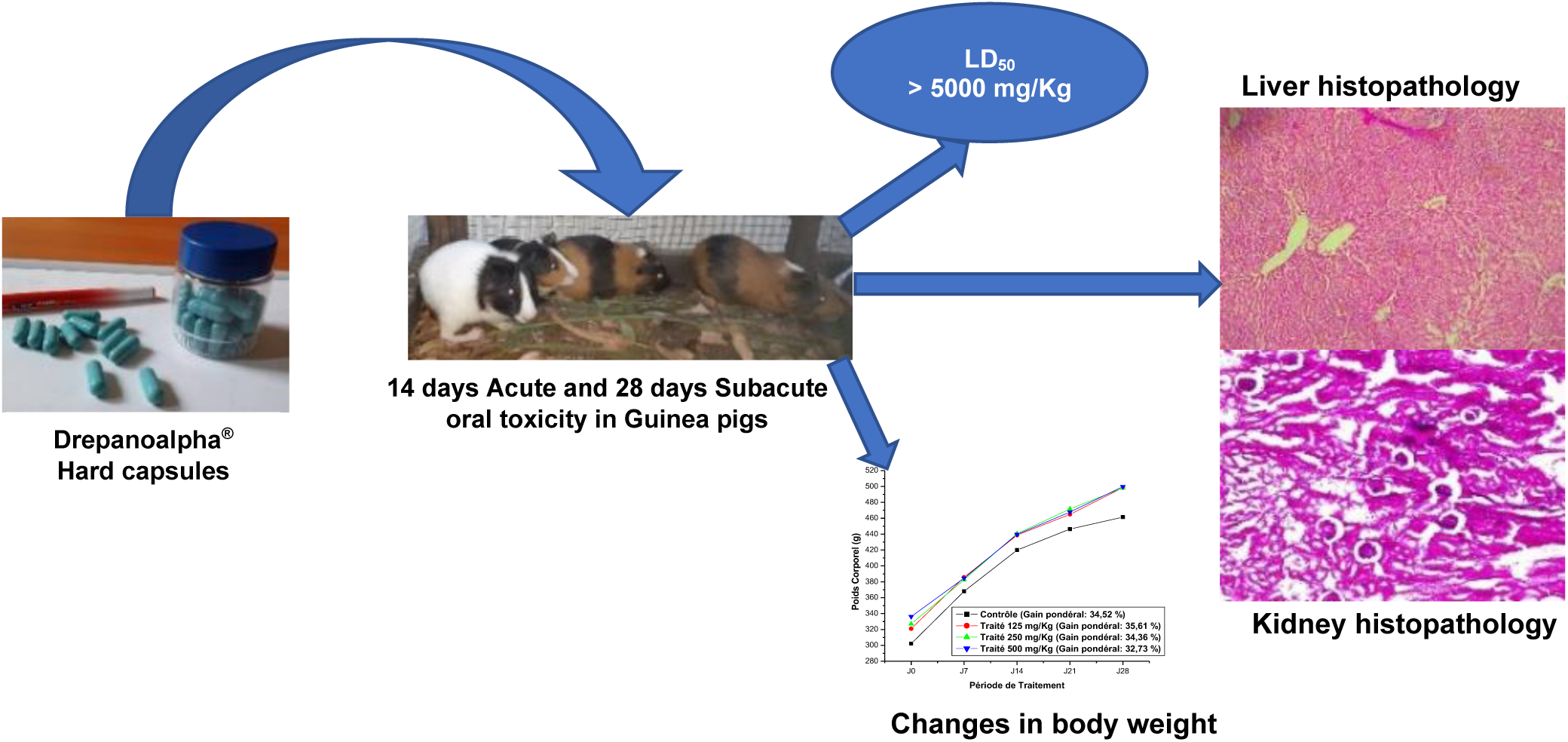

**ABSTRACT:** *Ethnopharmacological relevance:* Drepanoalpha^®^ hard capsules, a dry ethanolic extract (drug-extract ratio, 100/11) of a mixture of *Justicia secunda* Vahl and *Moringa oleifera* Lam dried leaves (1: 1, w/w) are used for the management of sickle cell disease in the Democratic Republic of Congo.

*Aim of the study:* This phytomedicine safety was investigated by acute and sub-acute administration in Guinea pigs.

*Materials and methods:* Healthy, male and nulliparous and non-pregnant female Guinea pigs were obtained from “Laboratory of Animal Experimentation and Toxicology” of the Department of Biology, Faculty of Sciences, University of Kinshasa. The animals were randomly selected, marked and divided into 2 groups of 5 animals each (3 males and 2 females) and 4 groups of 3 animals each for acute and sub-acute toxicity studies, respectively. The contents of hard capsules were dissolved in normal saline solution (NaCl 9 ‰). Animals received by gavage a single dose of 5000 mg/ kg of body weight (B.W.) of Drepanoalpha^®^ hard capsules (acute toxicity) and 125 mg/ kg, 250 mg/ kg and 500 mg/ kg of B.W. twice daily for 28 days (sub-acute toxicity). Normal saline solution was used as control. Hematological, biochemical and histopathological analyses were performed and the behavior of the animals was observed after treatment.

*Results:* The median lethal dose (LD_50_) is higher than 5000 mg/ kg of B.W., and the relative weights of vital organs (kidneys, liver and heart) collected from Guinea pigs at the end of treatment on D_14_ (acute toxicity) and D_28_ (sub-acute toxicity) has not undergone significant changes (p > 0.05). The results of haematological (red and white blood cells counts, haemoglobin, haematocrit) and biochemical (ALT, AST, albumin, total protein) tests did not show significant differences between control and test groups (α=0.05 for acute toxicity), while the histopathological study revealed no damage to the various organs excised.

*Conclusion:* The results indicate the safety of Drepanoalpha^®^ hard capsules, confirming previous studies, in rats and Guinea pigs, based on aqueous decoction of its raw herbs mixtures and the corresponding lyophilizate.

## 1. INTRODUCTION

Herbal medicines continue to gain popularity and interest in “green medicines” is increasing because they are considered to represent a supposedly safe alternative to modern medicine. Traditionally, plants and plant extracts are used to treat many diseases and disorders, but their use can represent serious health issues because of potential toxicities. Indeed, plant extracts can be very effective therapeutically, but some can reveal themselves to be toxic with a risk of serious poisoning (Chanda et *al*., 2015; Ngbolua et *al*., 2011a, b; Izzo, 2004; Whitto et *al*., 2003); moreover, mid-term to long-term toxicities represent particular risks as they are hardly detected by tradipraticians (Williamson et al., 2015). In the European Union, a safe use, correctly documented over a sufficient time, is considered to yield a plausible indication of innocuity, and forms the basis for marketing a traditional herbal product (Dickinson et al., 2019; HMPC, 2018). However, in many traditional medicines, and especially in developing countries, the use and safety of herbal products is hardly documented. Indeed, the use of medicinal plants as medicines is simply based on traditional and/or folklore uses that have often been perpetuated over several generations but not questioned, validated or documented (Mpiana et *al*., 2016; Sofowora, 1989). But, indeed, these products contain bioactive principles that may cause adverse effects (Bent and Ko, 2004). And so, for many plants with alleged therapeutic use, toxicity data are mandatory to ensure safety in the use of their medicinal products (Chanda et *al*., 2015).

According to the World Health Organization (WHO), more than 80 % of the population in poor regions of Africa mainly rely on traditional medicine (Mpiana et *al*., 2016; Chanda et *al*., 2015). Thus, in its 2014-2023 strategy, the WHO encourages African countries to develop and modernize Traditional African Medicine (TAM) as an integral part of their health care system (WHO, 2013). It has therefore endorsed the use of herbal products in national policies and drug regulatory measures to strengthen evaluation of their safety and efficacy (Mpiana et *al*., 2016).

The Democratic Republic of Congo (DRC) is home to nearly 47 % of Africa’s tropical forests and more than 80 % of its population depends on medicinal plants for the treatment of various diseases, including sickle cell disease, a genetic blood disorder resulting from a mutation of the ß-globin gene that causes the replacement of a glutamic acid residue with valine in the sixth position of the ß-chain of haemoglobin (Gbolo et *al*., 2017a; Ngbolua et *al*., 2013a).

Up to 2 % of the Congolese population is affected by this haemoglobinopathy (Thsilolo et *al*., 2009). Our research team has identified nearly 100 medicinal plants used by traditional healers to treat sickle cell disease (Mpiana et *al*., 2012). A bioguided selection of these plants led us to the formulation of Drepanoalpha^®^, that is a two-herbs formula produced from *Justicia secunda* Vahl and *Moringa oleifera* Lam dried leaves (Gbolo et *al*., 2017a; Mpiana et *al*., 2014a,b). Pre-clinical *in vitro* studies have reported that Drepanoalpha^®^ powder are endowed with antifalcemic, antihemolytic, antiradical and antioxidant properties (Gbolo et *al*., 2017b–c; Ngbolua et *al*., 2014d). Drepanoalpha^®^ powder have also shown an ability to increase red blood cell and platelets count as well as hemoglobin accumulation rate in rats, Guinea pigs and sickle cell patients and especially also its ability to suppress all crisis in sickle cell patients (Gbolo et *al*., 2017b-c; Ngbolua et *al*., 2014d); the mixture of herbs and their lyophilized extracts have no toxic effects on both rats and Guinea pigs (Mpiana et *al*., 2016; Ngbolua et *al*., 2014).

For more convenient administration and higher compliance, hard capsules of an extract were formulated and are currently under investigation. The content of capsules was investigated in the present study and designated as “Drepanoalpha^®^HC”

The overall objective of the present study is to evaluate in Guinea pigs the toxicity of “Drepanoalpha^®^HC”.

The specific objectives assigned to this research are:

- Evaluation of the LD_50_,
- Determination of haematological and biochemical parameters,
- Histopathological study of major organs (heart, stomach, liver and kidneys).
- Prediction of the ADMET (Absorption, Distribution, Metabolism, Excretion and Toxicity) profile of major compounds contained in “Drepanoalpha^®^HC”.

## 2. MATERIAL AND METHODS

### 2.1. Method of preparation and dose in traditional use

The phytomedicine Drepanoalpha^®^ powder, a 50-50 (w/w) mixture of two food plants, *Justicia secunda* Vahl Vahl (Herbarium MNHN-P-P00719831) and *Moringa oleifera* Lam (Herbarium MNHN-P-P05401821) leaves, is marketed in DR. Congo as a plant powder in 100 g plastic bags. For administration, a dose of Drepanoalpha^®^ powder, selected according to age (6 months to 2 years, 1/2 tsp; 3 to 8 years, 1 tsp; 9 to 10 years, 1.5 tsp; adults, 2 tsp), is infused in about 80 mL of boiling water, which is covered tightly and stirred from time to time for 20 min before filtering; 60 mL of this infusion is administered orally in 2 intakes (2 × 30 mL) per day.

### 2.2. Extraction and preparation of Drepanoalpha^®^ hard capsules

The dry extract (drug-extract ratio, DER, 100/11) was obtained by percolating the herbal material mixture with ethanol 96 %, and drying under vacuum at 40°C. The capsules were filled with a mixture of this ethanolic extract with expients, lactose and aerosil, in a ratio 12 : 80 : 8 (% w/w).

Figure 1 presents the HPTLC profile of the ethanolic extracts of Drepanoalpha^®^HC and its constituent plants.

**Figure 1.**
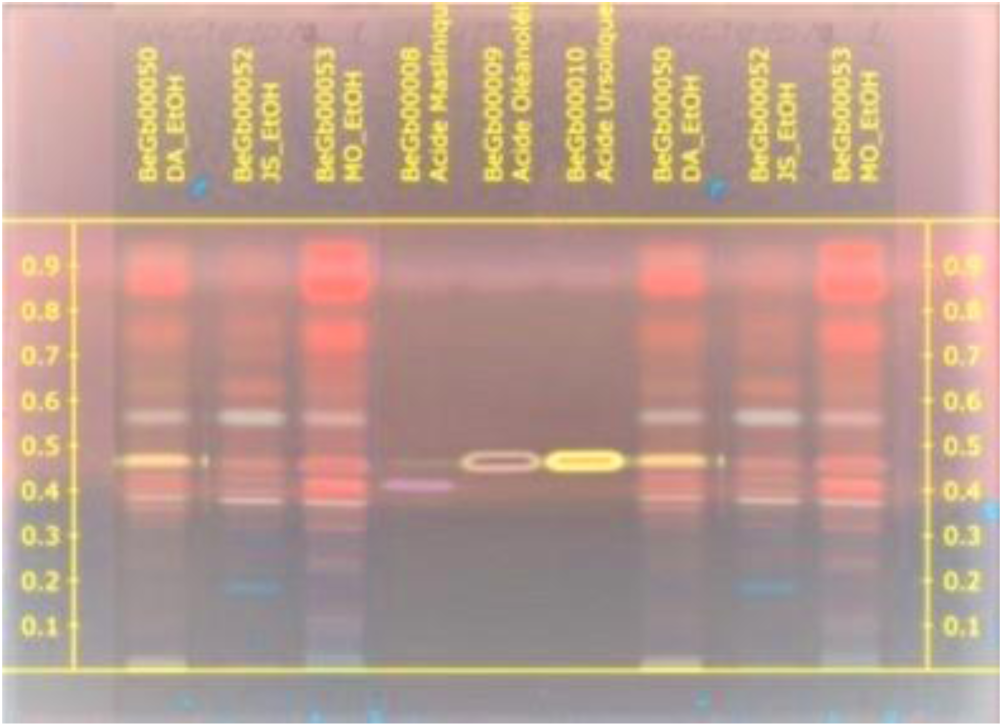
HPTLC profile of the ethanolic extracts of Drepanoalpha^®^HC and its constituent plants (5 mg/mL). Tracks: 1-Drepanoalpha^®^HC, 2-*Justicia secunda*, 3-*Moringa oleifera*, 4-maslinic acid, 5-oleanolic acid, 6-ursolic acid. Mobile phase: toluene - chloroform - methanol (60-20-20, V/V/V); derivatization with anisaldehyde-sulfuric acid; examination under UV_365nm_.

### 2.3. High performance thin-layer chromatography (HPTLC)

HPTLC was performed according to the procedure of the European Pharmacopeia 10 (COE, 2019), using Camag Automatic TLC Sampler (ATS 4), Automatic Developing Chamber 2 (ADC 2), Derivatizer and TLC Visualizer 2. The Camag systems were driven by the software visionCATS version 2.5. The HPTLC was performed on silica gel 60 F_254_ HPTLC plates (Merck, Germany); 2 µL of samples were applied in 8-mm wide bands, the plates were activated on MgCl_2_ (~ 33 % RH) and the tank saturated for 20 min; the solvent system was toluene - chloroform - methanol (60:20:20, V/V/V/V). The plate development was performed at 70 mm from the lower edge of the plate. After spraying an anisaldehyde-sulfuric acid mixture, derivatization was achieved by heating the plate at 105°C for 5 min and plates were visualized under UV_365nm_.

### 2.3. Selection and preparation of test animals

Healthy, male and nulliparous and non-pregnant female Guinea pigs were obtained from “Laboratory of Animal Experimentation and Toxicology” of the Department of Biology, Faculty of Sciences, University of Kinshasa. They were randomly selected, individually marked for identification purposes and acclimated to laboratory conditions for 3 days. Animal room temperature and relative humidity were 24 ± 2 ?C and 50–60%, respectively, and there was a 12 h light/dark cycle. Animals had free access to standard pelleted diet and fresh herbs, water ad libitum.

### 2.4. Protocols for toxicity studies

#### 2.4.1. Acute toxicity study (LD_50_)

Ten Guinea pigs were divided before fasting into two groups of 5 animals each (3 males and 2 females) according to their weight: a control group (mean weight: 305.03 ± 3.99 g) and a treatment group (weight: 326.14 ± 3.29 g). The test drug (prepared within the 1 h prior to administration) was administered by gavage in a single dose, using a gastric tube. All animals had free access to tap water and food throughout the experiment but were shortly fasted (withholding food, but not water) for 24 h before the oral administration of the single dose of drug. The control group received a normal saline solution (NaCl 9 ‰) and the treated group received a single dose of 5000 mg/kg of body weight of Drepanoalpha^®^ hard capsules content, suspended in the normal saline solution; the dose was selected according to the dose adjustment method of the Organization for Economic Cooperation and Development (OECD) guidelines 425 (OECD, 2008). Four hours after dosing, the animals had free access to water and food. Every day, the weight of all animals was measured.

#### 2.4.2. Sub-acute toxicity study

Twelve female Guinea pigs (OECD, 2008) were divided into 4 groups of 3 each. The administered doses were selected and distributed using arithmetic regression so that the highest administrated dose was equal to 1/10 of the LD_50_ value. The 1/10 of the LD_50_ for the case in point is the 1/10^th^ dose of the highest dose administered for acute toxicity. Thus, the experimental doses administered to the Guinea pigs of three groups were 125 mg/kg, 250 mg/kg and 500 mg/kg administered twice daily for 28 days, respectively, and the fourth group was the control group that received only a NaCl 9 ‰ solution.

#### 2.4.3. Signs of mortality and clinical signs (general behavior)

During the experimental period, all animals were observed daily for behavioral disturbances (difficult movement, drowsiness, slowed respiration and convulsions), latency and/or mortality were noted after dosing at 30 min, 1 hour, 2 hours, 5 hours, 24 hours and daily over 14 days for acute toxicity and 28 days for sub-acute toxicity.

As discussed in the OECD guideline (OECD, 2008), the LD_50_ is less than the test dose (5000 mg/kg) if at least three animals died and is higher than the test dose (5000 mg/kg) if at least three animals were able to survive.

#### 2.4.4. Measurement of body and organ weights

The body weight of each animal was measured with a sensitive scale (METTLER PM600) during the acclimatization period, once before the start of dosing and every day until the day of sacrifice. At the end of the experiment, all Guinea pigs were shortly fasted (but with free access to tap water) overnight from 6:00 p.m. to 6:00 a.m. and then were weighted. Three Guinea pigs per group were sacrificed by decapitation preceded by immobilization with ethanol.

The liver, kidneys, stomach and heart of each animal in the different groups were excised immediately after blood sampling. After excision, the organs were immediately weighed (paired organs were weighed together) and photographed and preserved with buffered formalin for histological examination. The organ weight ratio was calculated by the relationship:

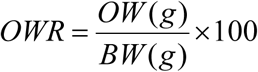

With OWR (organ weight ratio), OW (organ weight) and BW (body weight)

#### 2.4.5. Evaluation of histological damage

The liver, kidneys, stomach and heart of dissected animals were excised and preserved with buffered formalin for histopathological analysis according to the basic techniques described by Lee and Luna (1968). The individual excised organs were cut into several fragments, placed in cassettes and preserved in 10 % formalin. After inclusion in paraffin, the tissue blocks were sectioned in ribbons at a thickness of 4 *μ*m with a Leitz microtome (Leitz 1512). Finally, the organs slices were stained with haematoxylin-eosin, fixed between slide and coverslip before being observed using a microscope equipped with a Leica ICC50 W camera.

#### 2.4.6. Biochemical and haematological parameters

After decapitation, blood samples from each sacrificed animal were collected in EDTA (Ethylenediaminetetraacetic acid) tubes and non-EDTA dry tubes. EDTA-collected blood was used to determine hematological parameters; the red blood cells (RBC), hemoglobin (HGB), white blood cells (WBC) and hematocrit (HTC) were evaluated with an automated hematology analyzer (Coulter ABX Electronic). Serum was separated from non-EDTA-collected blood after centrifugation at 3000 rpm for 5 min and the following parameters were evaluated: alanine amino-transferase (ALT), aspartate amino-transferase (AST), urea, creatinine, total protein by spectrophotometric methods. The assays were carried out using the human brand biochemical automaton (HumaLyzer Primus, HUMAN, HUMAN Gesellschaft für Biochemica und Diagnostica mbH, Germany) with Biobase biochemical kits (HUMAN Gesellschaft für Biochemica und Diagnostica mbH, Germany).

### 2.5. ADMET Profile

Twelve molecules (Table 1) described in Drepanoalpha^®^’s plants (Kitadi et al., 2019; Ngbolua et al., 2018; Koffi et al., 2013) were selected according to their biological activities. These molecules include polyphenols (including flavonoids), fatty acid esters, triterpenoids, documented for their antisickling activities (Lukyamuzi et *al*., 2019; Tshilanda et *al*., 2016; Tshibangu et al, 2011), alkaloids known for their analgesic and anti-inflammatory activity (Hayfaa, 2013), thiocarbamates known for their anti-inflammatory and antitumor activities (Cheenpracha et *al*., 2010; Murakami, 1998). The pharmacokinetic properties (ADMET, i.e. absorption, distribution, metabolism, excretion and toxicity) of these twelve bioactive compounds were predicted *in silico*, using the pkCSM web server (http://biosig.unimelb.edu.au/pkcsm). The selected compounds were drawn using ChemDraw 14.0 software, and their SMILES were generated for calculations using the pkCSM web server.

**Table 1.** List of selected compounds

|  | Compound Name | Raw formula | Molecular weight | CAS number |
| --- | --- | --- | --- | --- |
| 1. | Niaziminin | C<sub>18</sub>H<sub>25</sub>NO<sub>7</sub>S | 399.46 g/mol | 147821-56-5 |
| 2. | Kaempferitrin | C<sub>27</sub>H<sub>30</sub>O<sub>14</sub> | 578.52 g/mol | 482-38-2 |
| 3. | Quinindoline | C<sub>15</sub>H<sub>10</sub>N<sub>2</sub> | 218.25 g/mol | 243-38-9 |
| 4. | Vasicine | C<sub>11</sub>H<sub>12</sub>N<sub>2</sub>O | 188.23 g/mol | 6159-55-3 |
| 5. | Ellagic acid | C<sub>14</sub>H<sub>6</sub>O<sub>8</sub> | 302.19 g/mol | 476-66-4 |
| 6. | 1,2,3-triolein | C<sub>57</sub>H<sub>104</sub>O<sub>6</sub> | 885.43 g/mol | 122-32-7 |
| 7. | 3,5,3'-trihydroxy-7,4'-dimethoxyflavone | C<sub>17</sub>H<sub>14</sub>O<sub>7</sub> | 330.29 g/mol | 529-40-8 |
| 8. | 3,5,7-trihydroxy-4'-methoxyflavone | C<sub>16</sub>H<sub>12</sub>O<sub>6</sub> | 300.26 g/mol | 57396-72-2 |
| 9. | 5,7-dihydroxy-3,4'-dimethoxyflavone | C<sub>17</sub>H<sub>14</sub>O<sub>6</sub> | 314.29 g/mol | 4712-12-3 |
| 10. | 5,4'-dihydroxy-7-methoxyflavanone | C<sub>16</sub>H<sub>14</sub>O<sub>5</sub> | 286.28 g/mol | 2957-21-3 |
| 11. | β-sitostérol | C<sub>29</sub>H<sub>50</sub>O | 414.71 g/mol | 83-46-5 |
| 12. | Stigmastérol | C<sub>29</sub>H<sub>48</sub>O | 412.69 g/mol | 83-48-7 |

Figure 2 presents the structures of selected molecules

**Figure 2.**
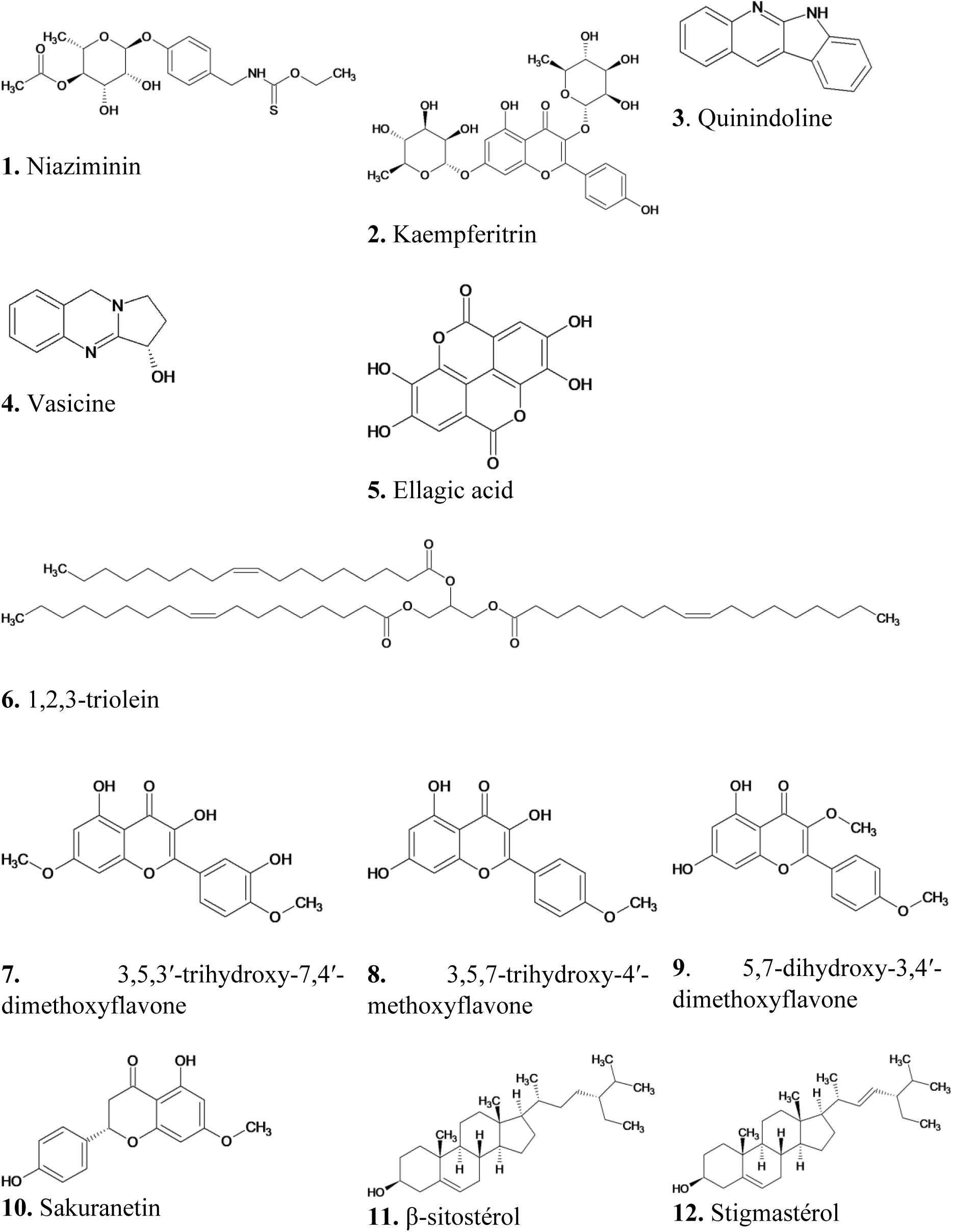
Structures of molecules **1-12** derived from Drepanoalpha^®^’s constituent plants

### 2.6. Statistical analysis

The statistical analyses of the results were performed using Origin version 6.1 software. The values are presented as an average ± SD. Significance levels between treatment and control groups were measured by the Student t-test and ANOVA. If p < 0.05, the difference between the values is considered significant.

### 2.7. Ethical considerations

The study was conducted after receiving the approval of the **Ethical and Scientific Committee of the School of Public Health** of the Faculty of Medicine of the University of Kinshasa, with the highest level of requirement for humanity and compassion in the use of animals in biomedical research (**Approval No.: ESP/CE/237/2019 of August 14, 2019**). The animals used in this study were not subjected to unnecessarily painful and frightening situations. The procedure was carried out by a well-trained person. The animals were protected from pathogens and placed in an appropriate environment. The number of animals was kept to the minimum possible, without compromising the scientific objectives of the study (Richmond, 2000, Veissier, 1999).

## 3. RESULTS

### 3.1. Acute toxicity study of Drepanoalpha^®^HC in Guinea pigs

The various signs of toxicity observed in **Guinea** pigs are described in **Table 2**.

**Table 2.**
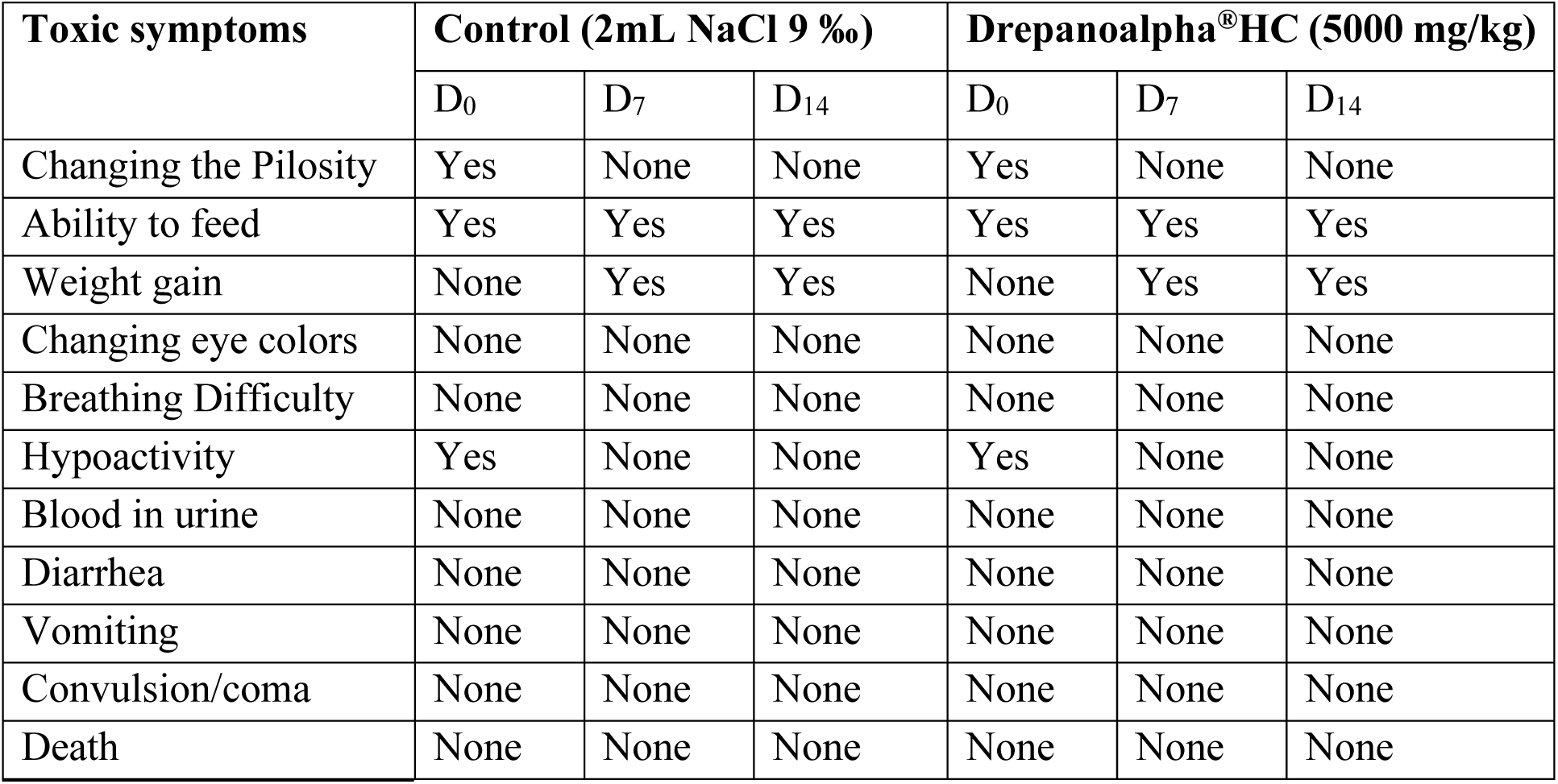
Acute toxicity of Drepanoalpha^®^HC administered by oral treatment.

This table shows that, during the acute toxicity observation period, none of the treated animals succumbed. No signs of toxicity such as tremor, change of step, blood in urine, convulsion, salivation, diarrhea, coma … were noted. However, a partial hypoactivity and a slight increase in the respiratory frequency in treated Guinea pigs was observed but these observations ended after about ten minutes. In addition to this hypoactivity and the transient change in respiratory rate, a modification of pilosity after a few minutes was also observed, which would reflect the stress felt during feeding. It took a few minutes for the animals to realize that they were not in any danger. These signs were identical in control and Drepanoalpha^®^HC-treated animals.

Table 3 presents the changes in body and organs weights of Guinea pigs after acute oral treatment with Drepanoalpha^®^HC

**Table 3.**
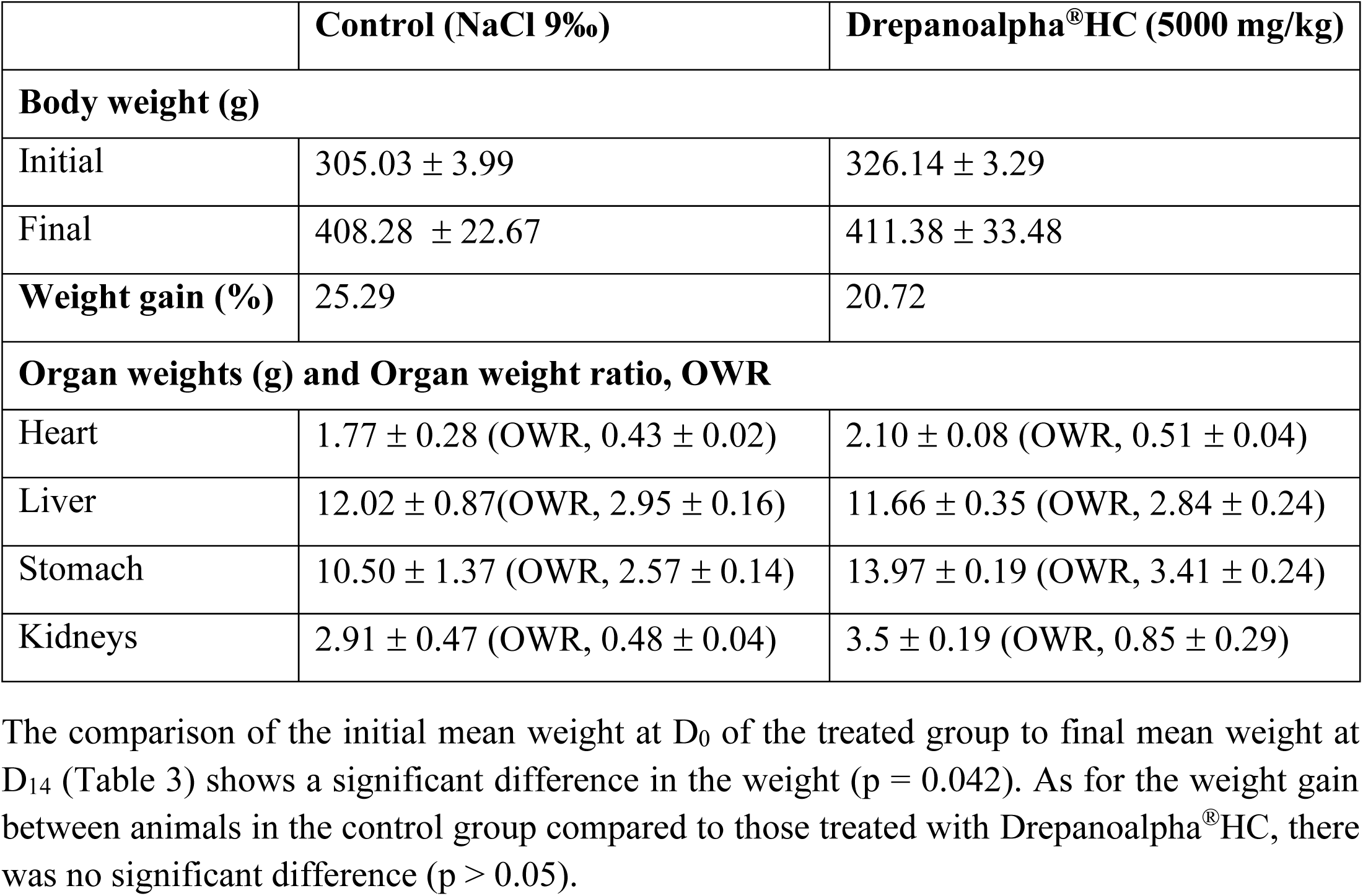
Changes in the body and organs weights of Guinea pigs after acute oral treatment with Drepanoalpha^®^ HC(n = 5)

The comparison of the initial mean weight at D_0_ of the treated group to final mean weight at D_14_ (Table 3) shows a significant difference in the weight (p = 0.042). As for the weight gain between animals in the control group compared to those treated with Drepanoalpha^®^HC, there was no significant difference (p > 0.05).

**Figure 3** displays the vital organs observed after dissection.

Statistical analysis applied to data presented in **Table 4** shows that on day D_14_ post-treatment, there was slight decrease (non-significant; p > 0.05) in WBCs, relatively to control.

**Table 4.** Effects of Drepanoalpha^®^HC on hematological parameters in acute toxicity study (on day 14).

| Haematological Parameters | Control (NaCl 9 ‰) | Drepanoalpa®HC (5000 mg/kg) |
| --- | --- | --- |
| Haemoglobin (g/dl) | 16.30 ± 0.30 | 16.70 ± 0.00 |
| White blood cells count (x10<sup>3</sup>/mm<sup>3</sup>) | 12.60 ± 4.70 | 11.60 ± 1.10 |
| Red blood cells count (x10<sup>6</sup>/mm<sup>3</sup>) | 4.10 ± 0.90 | 4.60 ± 0.00 |
| Haematocrit (%) | 49.00 ± 1.00 | 50.00 ± 0.00 |

**Figure 3.**
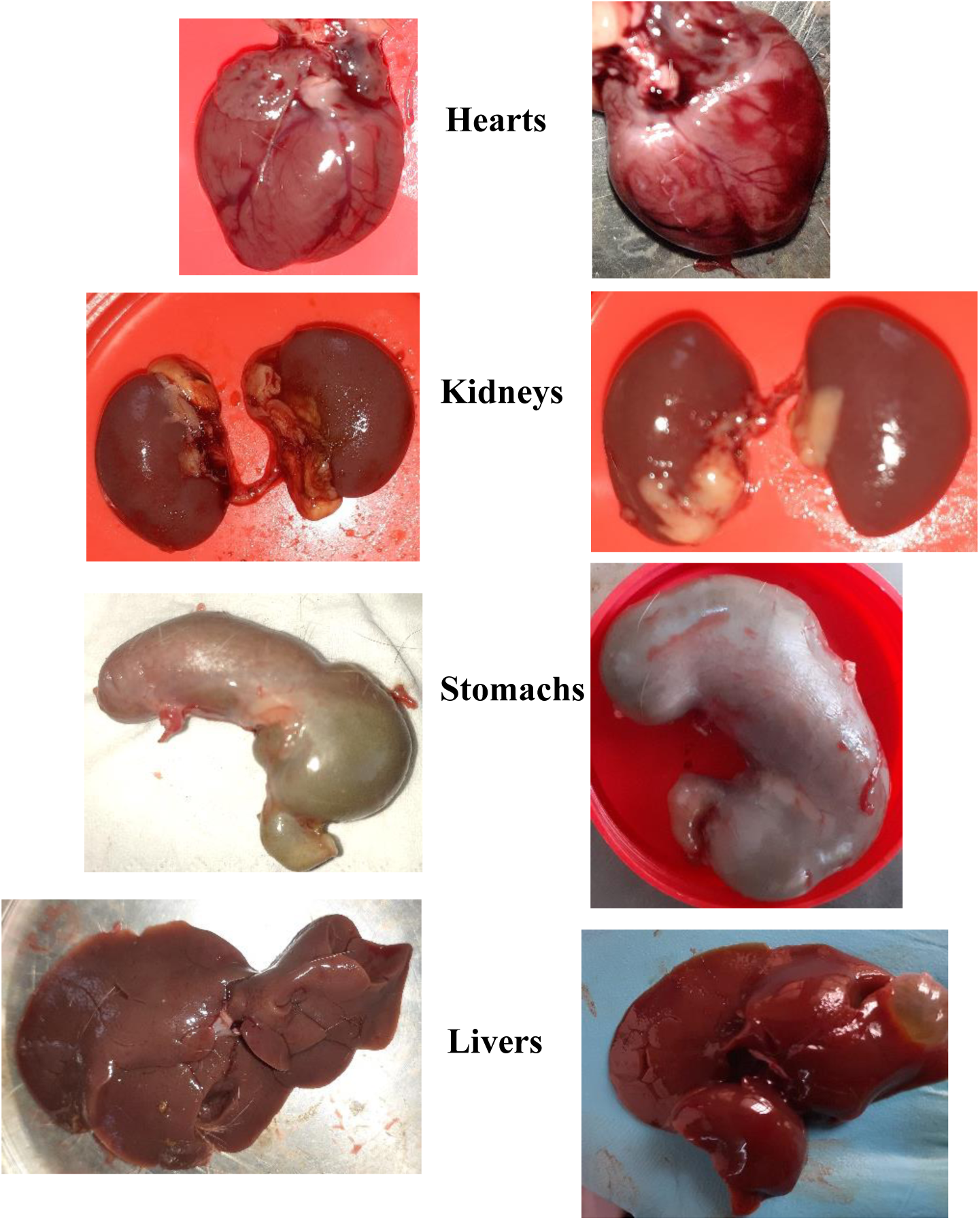
Photograph of vital organs of Control (NaCl 9 ‰; left) and Drepanoalpha^®^HC-treated (5000 mg/kg; right) Guinea pigs at day 14.

**Table 5** shows a small, but non significant, increase in serum AST and decrease in ALT in animals treated with Drepanolapha^®^. The transaminase level may vary, depending on parameters such as sex, age, body temperature, body mass index (BMI) and to the technique used in the medical analysis laboratory as well. Since the increase observed in the control group is not significant, we conclude that Drepanoalpha^®^HC is not toxic to liver cells.

**Table 5.** Effects of Drepanoalpha^®^HC on biochemical parameters in acute toxicity study (after 14 days).

| Biochemical | Control (NaCl 9‰) | Drepanoalpa® (5000 mg/kg) |
| --- | --- | --- |
| ALT (UI/l) | 72.10 ± 10.30 | 71.00 ± 11.80 |
| AST (UI/L) | 74.30 ± 33,50 | 78.20 ± 26.10 |
| Urea (g/L) | 0.20 ± 0.10 | 0.36 ± 0.10 |
| Creatinin (mg/L) | 7.30 ± 2.40 | 8.60 ± 2.50 |
| Direct bilirubin (mg/L) | 3.40 ± 0.40 | 0.700 ± 0.05 |
| Total bilirubin (mg/L) | 5.60 ± 0.40 | 3.70 ± 0.10 |

On day D_14_ post-treatment, no significant differences were observed between the mean values of creatinine and urea for the two groups. These results indicate that Drepanoalpha^®^HC is not toxic to the kidneys.

**Table 6** shows that Drepanoalpha^®^HC does not disturb the mean values of total protein and albumin on day D_14_ after treatment (p > 0.05). This is experimental evidence that the biosynthetic capacity of the liver is maintained intact and therefore the product is not hepatotoxic.

**Table 6.**
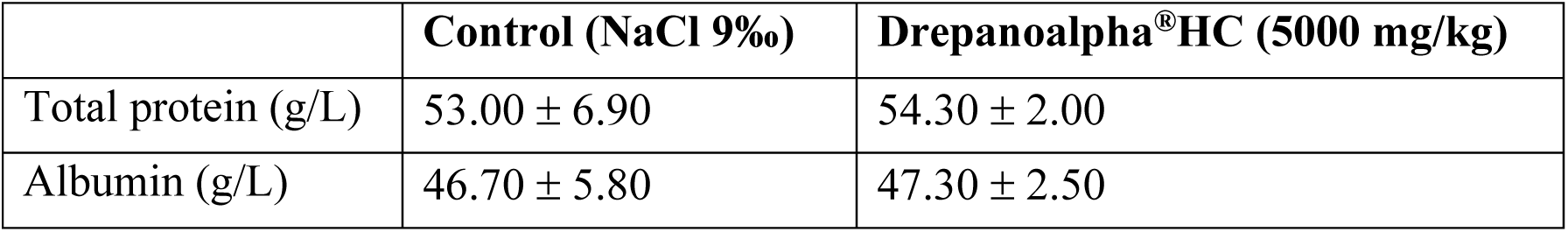
Effects of Drepanoalpha^®^HC on blood proteins in acute toxicity study (after 14 days).

Histological analysis of different organs of Guinea pigs treated with Drepanoalpha^®^HC (Figure 4) shows a normal morphology in all rodents.

**Figure 4.**
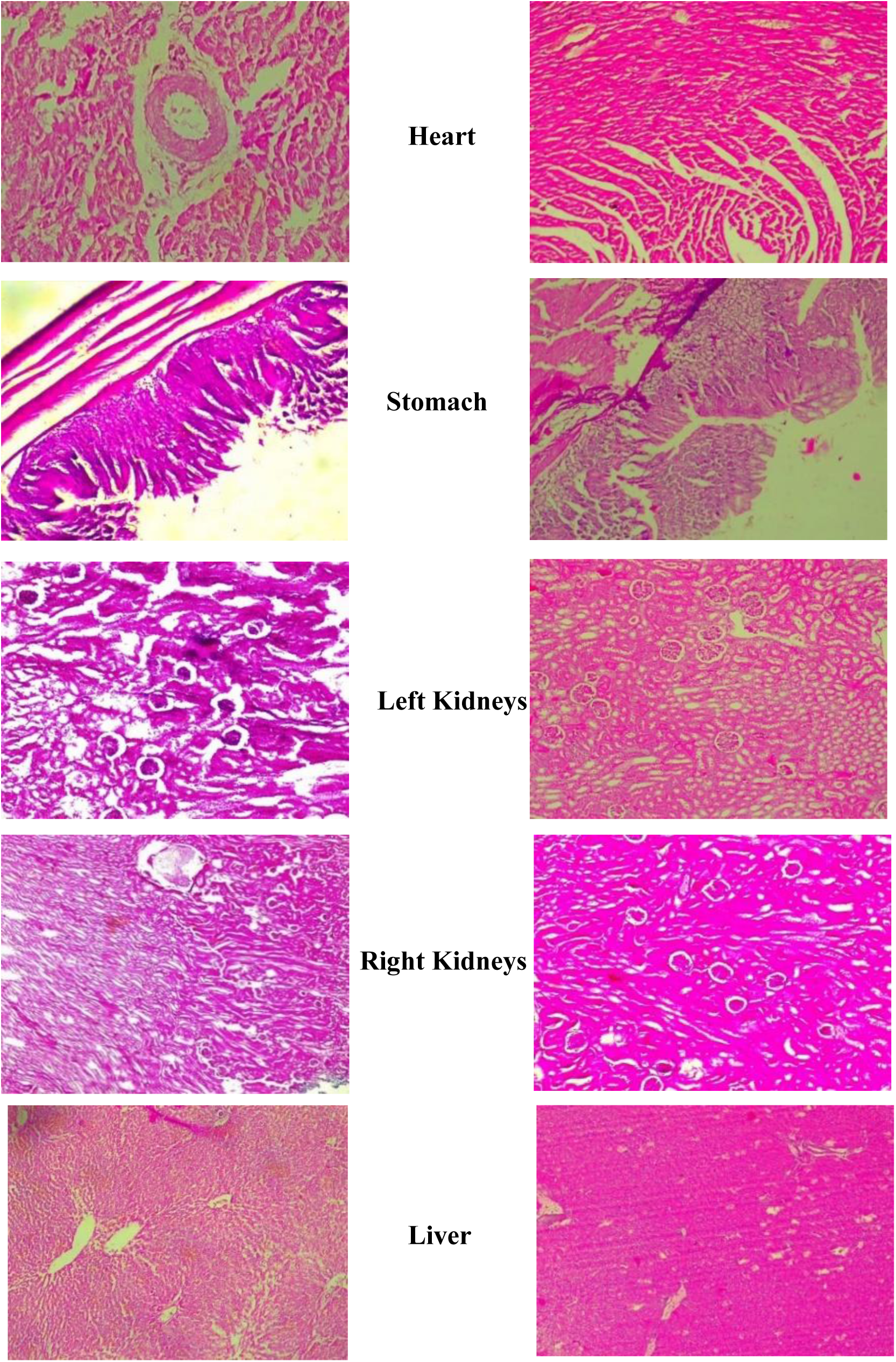
Comparison of histological sections between Control (NaCl 9 ‰; left) and Drepanoalpha^®^HC-treated (5000 mg/kg; right) Guinea pigs at day 14.

### 3.2. Sub-acute toxicity study of Drepanoalpha^®^HC in Guinea pigs

**Figure 5** shows the effects of Drepanoalpha^®^HC on body weight (g) and weight gains of Guinea pigs in the sub-acute toxicity study.

**Figure 5.**
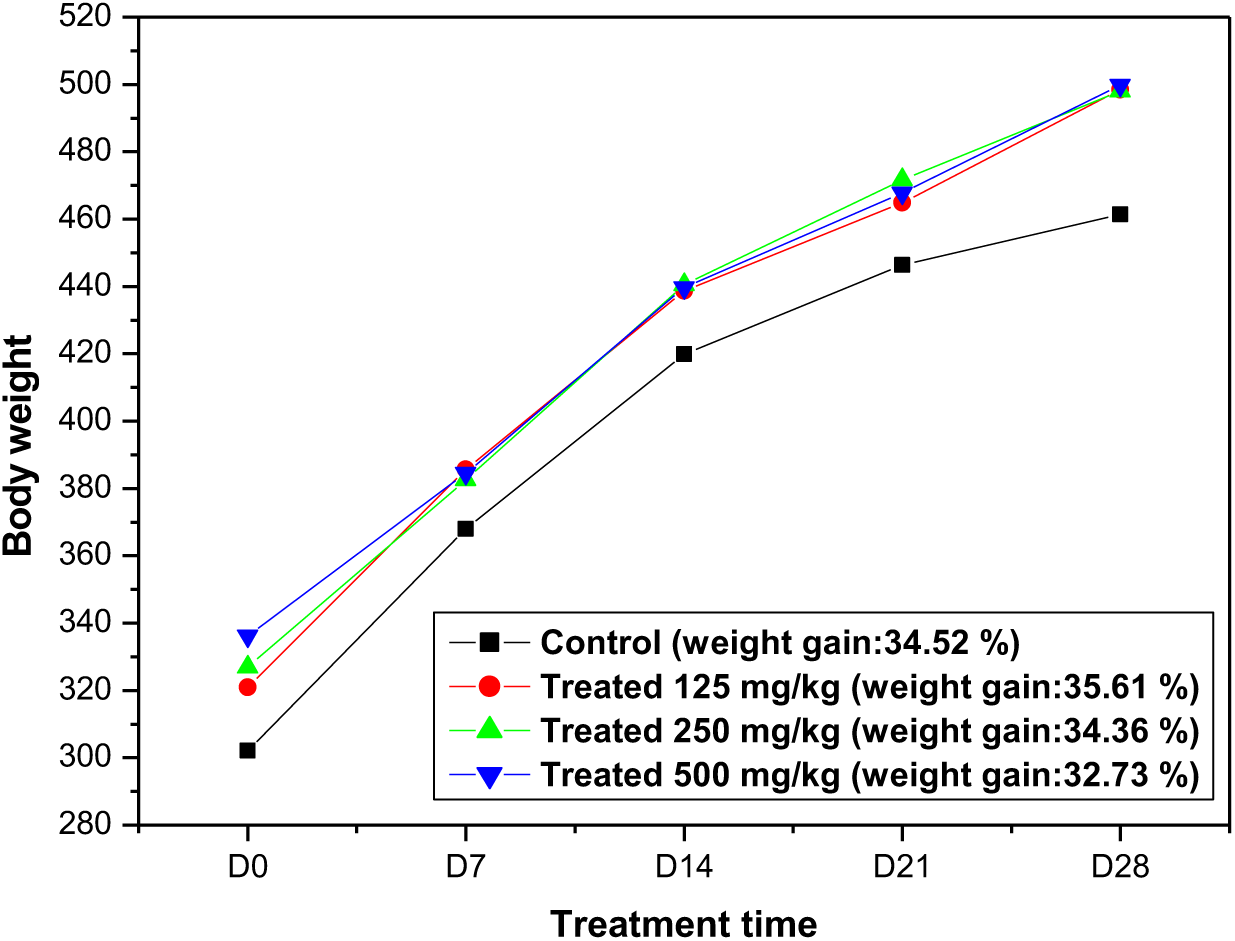
Effect of Drepanoalpha^®^HC on guinea pig body weight (g) and weight gain (%) in the subacute toxicity study (28 days).

Comparisons of the initial and final mean weights of Guinea pigs of the control and treated groups on D_28_ (**Figure 5**) show no significant differences. Accordingly, the weight gains between animals in the control group and those in the Drepanoalpha^®^HC-treated groups at different doses for 28 days show no statically significant difference (p > 0.05).

**Table 7.**
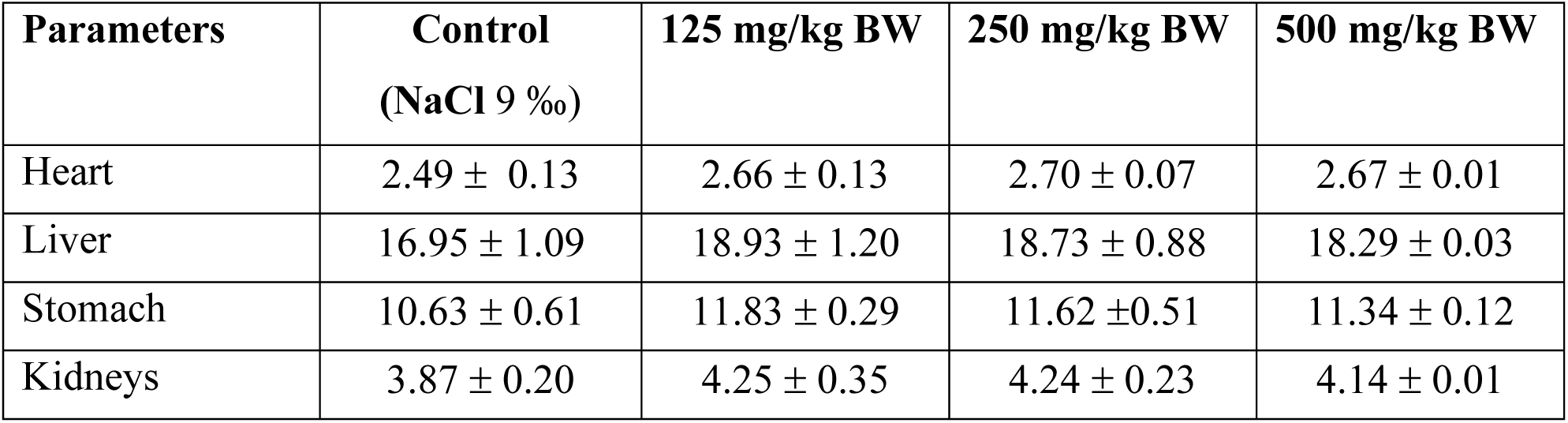
Effect of Drepanoalpha^®^HC on organ weights (g) of Guinea pigs in sub-acute toxicity study (28 days).

The 28-days sub-acute toxicity oral administration of Drepanoalpha^®^HC in Guinea pigs induced no adverse effects on target vital organ weights. The comparison of organ weights of guinea pigs treated with Drepanoalpha^®^ at different concentrations with those of the control group showed no statically significant difference (p>0.05).

**Table 8.** Effect of different doses of Drepanoalpha^®^HC on hematological parameters in sub-acute toxicity study (28 days).

| Parameters | Control (NaCl 9‰) | Drepanoalpha<sup>®</sup>HC 125 mg/ kg | Drepanoalpha<sup>®</sup>HC 250 mg/kg | Drepanoalpha<sup>®</sup>HC 500 mg/kg | P for the difference between groups |
| --- | --- | --- | --- | --- | --- |
| Haemoglobin (g/dl) | 13.23 ± 0.51 | 14.23 ± 1.07 | 14.43 ± 0.84 | 15.33 ± 0.67 | 0.012 |
| White blood cells count ($\times 10^3/\text{mm}^3$) | 16.62 ± 0.16 | 12.10 ± 0.66 | 10.83 ± 1.53 | 11.50 ± 1.50 | 0.004 |
| Red blood cells count ($\times 10^6/\text{mm}^3$) | 4.75 ± 0.22 | 5.42 ± 0.21 | 5.74 ± 0.30 | 6.37 ± 0.25 | 0.001 |
| Haematocrit (%) | 39.67 ± 1.53 | 42.67 ± 3.21 | 43.33 ± 2.52 | 46.00 ± 2.00 | 0.012 |

The 28-days oral sub-acute toxicity administration of Drepanoalpha^®^HC in Guinea pigs shows a statistically significant (p > 0.05) decrease in WBC counts and increase in the other parameters, red blood cell count, hemoglobin, and hematocrit.

**Table 9.** Effect of different doses of Drepanoalpha^®^HC on biochemical parameters in sub-acute toxicity (28 days).

| Parameters | Control (NaCl 9 ‰) | Drepanoalpha<sup>®</sup>HC 125 mg/ kg | Drepanoalpha<sup>®</sup>HC 250 mg/kg | Drepanoalpha<sup>®</sup>HC 500 mg/kg |
| --- | --- | --- | --- | --- |
| ALT (UI/l) | 60.63 ± 10.35 | 53.35 ± 7.36 | 47.28 ± 12.05 | 41.63 ± 3.15 |
| AST (UI/L) | 32.52 ± 7.30 | 24.10 ± 3.24 | 25.95 ± 5.64 | 22.81 ± 6.01 |
| Urea (g/L) | 27.37 ± 9.48 | 23.58 ± 3.25 | 24.07 ± 5.32 | 23.48 ± 9.22 |
| Creatinin | 8.10 ± 1.20 | 6.70 ± 0.40 | 6.17 ± 0.20 | 6.17 ± 0.20 |
| Direct Bil. | 3.07 ± 2.20 | 2.47 ± 0.60 | 2.53 ± 0.40 | 2.67 ± 0.80 |
| Total Bil. | 5.77 ± 2.10 | 3.67 ± 1.60 | 4.20 ± 1.10 | 4.23 ± 0.30 |

Biochemical analysis of the 28-day oral sub-acute toxicity test in guinea pigs showed a non-significant decrease in all parameters at the different doses compared to the control. Only the ALT values at 500 mg/kg of body weight showed a significant reduction (p = 0.038).

**Table 10.** Effect of different doses of Drepanoalpha^®^HC on blood proteins in sub-acute toxicity study (28 days).

| Parameters | Control (NaCl 9 ‰) | Drepanoalpha<sup>®</sup>HC<br>125 mg/ kg | Drepanoalpha<sup>®</sup>HC<br>250 mg/kg | Drepanoalpha<sup>®</sup>HC<br>500 mg/kg |
| --- | --- | --- | --- | --- |
| Total protein | 53.00 ± 6.90 | 54.33 ± 2.10 | 48.33 ± 2.50 | 46.00 ± 4.40 |
| Albumin (g/L) | 38.00 ± 1.00 | 29.67 ± 3.80 | 31.33 ± 2.10 | 29.33 ± 5.90 |

Drepanoalpha^®^HC does not interfere with the mean values of total protein and albumin on day D_28_ post-treatment in oral sub-acute treatment (p > 0.05).

**Table 11.**
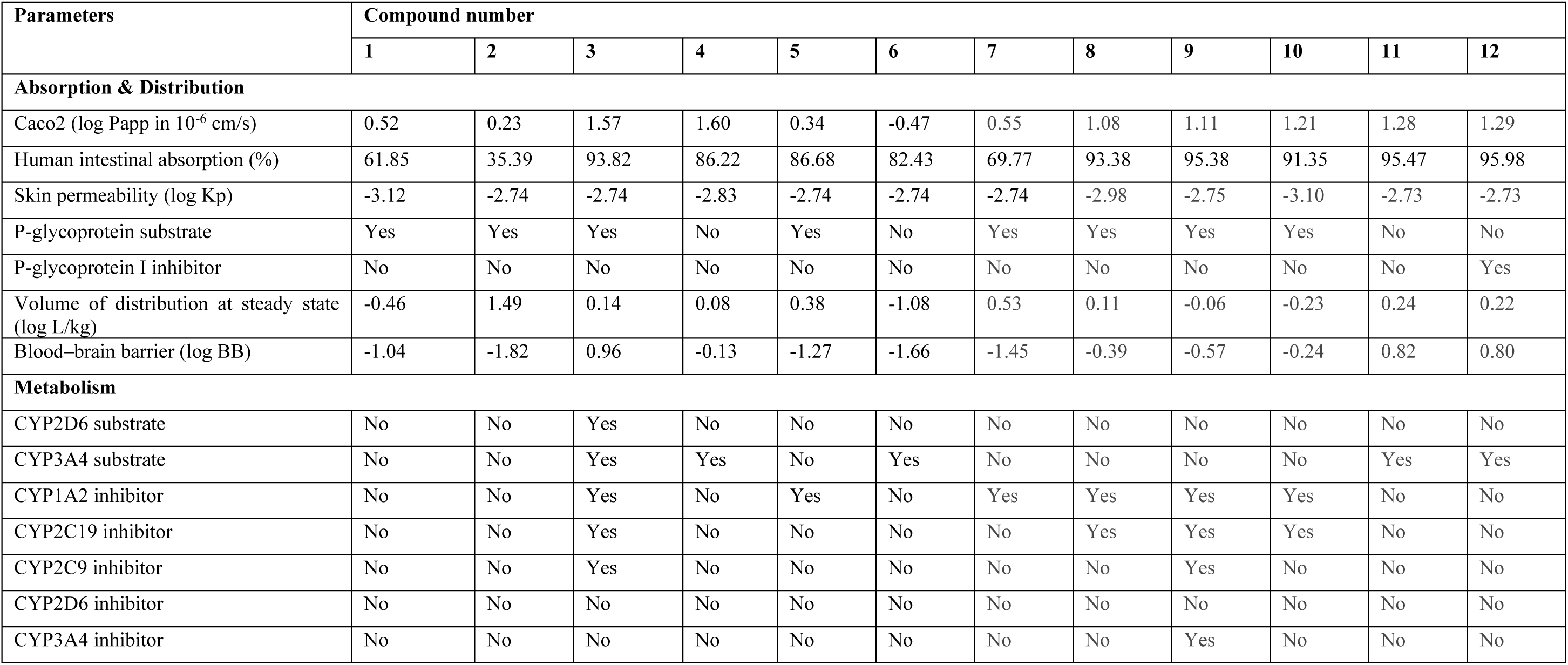
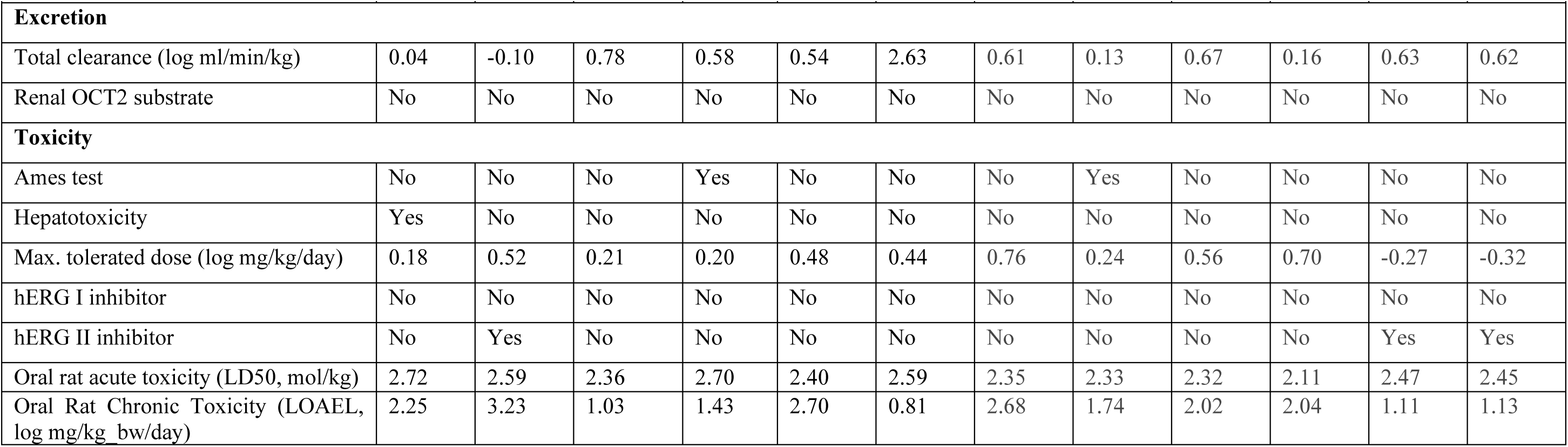
ADMET profiles of molecules **1-12**

## DISCUSSION

Phytotherapy is used by up to 80 % of the population in developing countries. Despite its widespread use, few scienti?c studies have been undertaken to verify the safety and efficacy of traditional remedies (Hilalya et *al*., 2004). To assess the safety of Drepanoalpha^®^ hard capsules, an improved traditional medicine for human use, toxicological evaluation in experimental models allowed to provide indications for the selection of a minimum tolerated dose for humans. The choice of rodents for this purpose is related to the fact that these animals express side effects similar to humans in terms of hematological, gastrointestinal and cardiovascular effects (Chukwuma and Obioma, 2014; Ogbonnia et *al*., 2010).

### Acute toxicity study

The acute toxicity test, the first step in the toxicological evaluation of drugs, consists in treating animals with a single dose within a short period of time and observing the adverse effects and mortality of the animal (Joshua et *al*., 2008). In this study, Drepanoalpha^®^HC induced no mortality at the limit dose of 5000 mg/kg of B.W. during the observation period of 14 days. However, slight changes in behavior, respiration and hypoactivity were observed in treated Guinea pigs for a few minutes after dosing. This transient hypoactivity was also present in control animals, indicating a slight trauma of administration conditions. The animals need a resting time necessary for the absorption of the extract (Ngbolua et *al*., 2016). Such slight behavioral deviations would be due to an adaptation following ingestion of a product that is not part of their diet. Regarding natural products, the observation of a respiratory frequency reduction, as well as convulsions could be observed upon intoxication with phenolic compounds; whereas hypoactivity could be attributed among others to sedative, anaesthetic and especially psychoactive properties of compounds, notably alkaloids (Luis Blanc et *al*., 1988 in Lohoues et *al*., 2006). Although such secondary metabolites are present in Drepanoalpha^®^, either they are non toxic by themselves or present at non-toxic dosages. The test limit for acute oral toxicity is generally considered to be 5 g/kg of body weight. If no mortality is observed at this dose level, a higher dose is generally not required (Hayes, 2011; OECD, 2008). The absence of mortality at this dose level indicates that there are no phytomedication-related adverse effects and that Drepanoalpha^®^ is not toxic. These results are in agreement with those obtained by Ngbolua et al (2016) and Mpiana et al (2014) who worked on the aqueous decoction of Drepanoalpha^®^ plant powder and its lyophilisate, respectively. Consequently, these results suggest that the median lethal dose (LD_50_) is higher than 5000 mg/kg. These results show that Drepanoalpha^®^, i.e. the plants ethanolic extract and excipients (lactose and aerosil) mixture, is practically non-toxic or non-lethal after acute exposure in guinea pigs. According to Kennedy et al (1986) and Diezi (1989), substances with an oral LD_50_ greater than 5 g/kg can be considered practically non-toxic; furthermore, according to the Hodge and Sterner (1980) scale, Drepanoalpha^®^ has a toxicity index of 5, thus almost non-toxic, suggesting safety (Kotué et *al*., 2013; Diezi, 1989).

Changes in body weight are generally used as an indicator of the side effects of drugs and chemicals compounds (Tofovic and Jackson, 1999; Raza et *al*., 2002; Teo et *al*., 2002; Hilalya et *al*., 2004; Nandy et *al*., 2012; Kingsley et *al*., 2012), any weight gain indicates that the drug does not alter carbohydrate, lipid and protein metabolism. The increase we observe in body weight of test animals suggests that they have increasingly accumulated calories from their normal diet (Chukwuma and Obioma, 2014). Although the diet of experimental animals is already rich in nutrients, it was expected that Drepanoalpha^®^ may have allowed a higher absorption and use of these nutrients as its constituent plants contain some trace elements (Gbolo et *al*., 2017); however there was no difference between control and treated groups. This no difference in treated groups is nevertheless further evidence that Drepanoalpha^®^ is safe. Indeed, a toxic extract generally leads to a weight loss of about 10 % or more (Schilter et *al*., 2003, Chanda, 2015). These results corroborate those obtained previously on and Drepanoalpha^®^ powder and its lyophilisate (Mpiana et *al*., 2016; Ngbolua et *al*., 2014).

Organs weight is also an important indicator of the physiological and pathological status of the animal. It is well established that organ weights vary with age and/or body weight. Indeed, it makes it possible to demonstrate whether or not an organ is attacked upon drug exposure (Mukinda and Syce, 2007; Ramesh et *al*., 2009). Heart, stomach, liver and kidneys are organs that can be damaged by an intoxication. Especially susceptible are kidneys, organs implied in major excretion pathways, and liver, the key organ in the metabolism and detoxification of xenobiotics (Clarke and Clarke, 1977; Viau and Tardif, 2003; Ramesh et *al*., 2009); a difference in weight could be attributed to a high rate of excretion and/or metabolism following exposure to some secondary metabolites (Jothy et *al*., 2001). The heart, can be affected by chemical compounds (Oduola et *al*., 2010), resulting in either gross or more subtle (arrythmia) effects. In this study, no macroscopic signs of toxicity (no mortality and organ damage) were observed during the study period on the tested organs (heart, stomach, kidneys, liver). Hematological parameters are also a good indication in the toxicological risk assessment of drugs. Indeed, blood constituents provide information on toxic effects because the hematopoietic system is one of the highly sensitive targets and an important indicator of physiological and pathological status, in both humans and animals (Jothy et *al*., 2001, Adeneye, 2006; Mukinda and Syce, 2007, Chanda, 2015). The slight, but not significant, increase in parameters WBC, HCT and HB, observed in Guinea pigs on day 14 following massive ingestion of Drepanoalpha^®^ can be correlated with the observation discussed hereunder for sub-acute treatments.

Biochemical parameters are involved in the determination of overall toxicity, indicating lesions to specific organs or systems; urea/creatinin and transaminases are abnormally elevated in the blood when drugs are toxic to kidneys and liver, respectively (Chukwuma and Obioma, 2014; Ngbolua et *al*., 2014). When the kidneys are damaged, less urea is eliminated and therefore remains in the bloodstream, where it accumulates; creatinin is a good indicator of the glomerular filtration function and high levels of creatinin in the blood correlate with poor kidney function (Gnanamani et *al*., 2008). In this study, no alterations in the values of these two parameters were observed. In case of hepatocyte lysis, cytosolic enzymes are released into the blood; the serum activity of enzymes ALT (GPT) and AST (GOT) is a conventional means of evaluating the functional integrity of the liver (Aliyu et *al*., 2006; Gome et *al*., 2011; Dufour et *al*., 2000; Pratt and Kaplan, 2000; AlHabori et *al*., 2002). ALT and AST, intracellular enzymes involved in amino acid metabolism, are highly concentrated in muscle, liver and brain; an increased concentration/activity of these enzymes in the serum is an indication of necrosis to these tissues (Hussein and Abdel-Gawas, 2010; Forchielli et *al*., 2012). ALT and AST levels rise rapidly when the liver is damaged, for example in hepatic cell necrosis, hepatitis or cirrhosis, associated with hepatotoxicity or viral attack (Pratt and Kaplan, 2000; Dufour et *al*., 2000). In the case of myocardial infarction, AST usually increases by 10 to 20 times the normal mean, ALT being normal or only slightly increased (Iwuanyanwu et *al*., 2012). Our results indicate that Drepanoalpha^®^HC did not significantly alter these parameters. Serum proteins are part of a general health check-up, to determine nutritional status, or to look for certain liver or kidney disorders. Albumin (synthesized by hepatocytes) represents 60 % of the proteins present in the blood, transporting many endogenous and exogenous substances in the blood and helping to maintain oncotic pressure. High albumin levels in the blood can be the result of hemoconcentration due to dehydration, fluid loss or diabetes insipidus. But it will also indicate poor transport of substances (Viau and Tardif, 2003). While a low albumin level in the blood reflects undernutrition, related to anorexia nervosa or tumours, hepato-cellular insufficiency, various severe inflammations or kidney damage. This study, indicates that Drepanoalpha^®^HC does not alter serum protein values.

Histopathological study of the kidney, liver, stomach and heart recovered from treated and control animals showed normal architectures (Figure 3), suggesting that no detrimental changes and morphological disturbances were caused by the single oral administration of Drepanoalpha^®^. These results corroborate those of the biochemical analyses on kidney and liver function.

### Subacute toxicity study

After 28 days of subacute toxicity asministration and observation, no signs of intoxication were noted for the groups treated with Drepanoalpha^®^HC at the 3 experimental dose, i.e. twice daily 125 mg/kg, 250 mg/kg and 500 mg/kg. Weight gain was also observed in all guinea pigs, similarly to those of the control group A statistically significant (p > 0.05) decrease in WBC counts and increase in the other parameters, red blood cell count, hemoglobin, and hematocrit could be explained by an increased production of haematopoietic regulatory factors (Kafaie et *al*., 2012). These observations show that Drepanoalpha^®^HC could prevent painful attacks in sickle cell disease patients and are in perfect agreement with our previous work on Drepanoalpha^®^ (Mpiana et *al*., 2016; Ngbolua et *al*., 2014; Gbolo et *al*., 2017). These results on sub-acute toxicity confirm those of our previous work on acute and sub-acute oral toxicity study of Drepanoalpha^®^ powder (Mpiana et *al*., 2016).

### In silico ADMET study on major compounds of Drepanoalpha^®^

The metabolism of xenobiotics is made possible by a series of enzymes, notably cytochromes P450 (CYP) that play a central role in their biotransformation, metabolism and/or detoxification. The latter are divided into families (CYP 1-2-3) and subfamilies (CYP 1A - 2C - 2D - 3A), the main ones involved in drug metabolism being CYP1A2, CYP2C8, CYP2C9, CYP2C19, CYP2D6, CYP3A4. Drug metabolism most often involves several CYPs and more rarely a single or preferential CYP: it is under these circumstances that the risk of interaction is highest (Guéguen et al., 2006).

*In vivo* toxicological studies (acute and sub-acute toxicity) in rodents and humans have shown that Drepanoalpha^®^ is not toxic. However, in order to prevent drug interactions, twelve molecules isolated from the constituent plants of this phytomedicine were analyzed *in silico* for their pharmacokinetic and toxicological properties (ADMET).

Eight of these compounds (1, 2, 3, 5, 7, 8, 9 and 10) are substrates for the P-glycoprotein. The metabolism of xenobiotics breaks down into 3 phases (functionalization: oxidation-reduction and hydrolysis, conjugation and transportation) that ultimately lead to the elimination of foreign substances in bile and urine. Glycoprotein P (P-gP) is involved in the intestinal absorption, passage through the foeto-maternal barrier and biliary and kindey transports, and especially of conjugated derivatives (Guéguen et *al*., 2006).

Compound 3 is an inhibitor of cytochromes CYP1A2, CYP2C19 and CYP2C29. Compound 9 is an inhibitor of CYP1A2, CYP2C19, CYP2C29 and CYP3A4, while compounds 5, 7, 8 and 10 are inhibitors of CYP1A2 and compounds 8 and 10 are also inhibitors of CYP2C19.

These results indicate that the presence of some of these compounds in the phyotomedine can lead to interactions with conventional drugs, transoported by P-gP and/or metabolized by CYP1A2, CYP2C19, CYP2C9 and/or CYP3A4. Such interactions have been reported to extend the range of plamatic concentrations variability to about 400-fold. However, these differences in metabolism, associated with plasma drug concentrations, do not translate into linear differences in drug activity, effects depending on the relationship between drug concentration, receptor/enzyme coupling efficiency, i.e. tissue-dependent potency, efficacy, and adverse effects (Wilkinson, 2005).

In addition, at high concentrations, compound 1 is predicted to cause hepatoxicity, compounds 2, 11 and 12 cardiotoxicity while compounds 4 and 8 could be genotoxic. At therapeutic dosages, our data indicate that the Drepanoalpha^®^ tested here would be non-toxic; this shows the importance of standardization in eventually detrimental compounds and strict adherence to the dosage. Indeed, the expression of these compounds in the plant is a function of *(i)* abiotic factors such as climate, geological environment of the plant harvesting site, etc.; *(ii)* biotic factors such as the presence of predators and/or parasites as well as interspecific competitions within ecosystems; and *(iii)* genetic factors, the specificity of each plant taxon (species, variety, cultivar) being related to its genetic constitution (Ngbolua, 2012).

## CONCLUSION

Acute (5000 mg/kg) and sub-acute (2 × 125, 250 and 500 mg/kg for 28 days) oral toxicity studies of Drepanoalpha^®^ in Guinea pigs showed no detrimental effects nor mortality, either during treatment or the observation period. These results suggest that oral administration of this formulation does not induce any toxic effects and that Drepanoalpha^®^ ethanol extracts are safe.

## Contributions of the authors

Gbolo B.Z. designed and carried out the study, conducted the experiments, analysed the data and wrote the manuscript. Mpiana P.T., Duez P., Ngbolua K.N., Tshibangu D.S.T., Memvanga P. and Tshilanda D.D. helped design the study and supervised it. Lebwaze Bienvenu assisted with the histopathological study. Aristote Matondo and Kilembe J.T. helped to carry out calculations on ADME-T properties. NACHTERGAEL A. helped to design and provide information for all the molecules. All authors read and approved the final manuscript.

## Acknowledgements

This project was carried out thanks to the funding of the Academy of Research and Higher Education (ARES) in its programm “Institutional Support (IA) 2018-2020”. The authors would like to thank the Laboratory of Pathological Anatomy of the University of Kinshasa (DRC) for the realization of histopathological sections; the Laboratory of Biochemical Hematology for the hematological and biochemical analyses. To Gabriel Moke and Misengabu Nicole for their technical assistance.

